# Genomic and protein structure modelling analysis depicts the origin and infectivity of 2019-nCoV, a new coronavirus which caused a pneumonia outbreak in Wuhan, China

**DOI:** 10.1101/2020.01.20.913368

**Authors:** Ning Dong, Xuemei Yang, Lianwei Ye, Kaichao Chen, Edward Wai-Chi Chan, Mengsu Yang, Sheng Chen

## Abstract

Detailed genomic and structure-based analysis of a new coronavirus, namely 2019-nCoV, showed that the new virus is a new type of bat coronavirus and is genetically fairly distant from the human SARS coronavirus. Structure analysis of the spike (S) protein of this new virus showed that its S protein only binds weakly to the ACE2 receptor on human cells whereas the human SARS coronavirus exhibits strongly affinity to the ACE receptor. These findings suggest that the new virus does not readily transmit between humans and should theoretically not able to cause very serious human infection. These data are important to guide design of infection control policy and inform the public on the nature of threat imposed by 2019-nCov when results of direct laboratory tests on this virus are not expected to be available in the near future.

---

A cluster of pneumonia cases of unknown cause were reported in Wuhan, the capital City of Hubei Province of China in December 2019. As of 18^th^ January 2020, a total of 44 such cases were documented, among whom two patients have died, five in critical condition, and six have been discharged from hospital. Most patients had visited or worked in a seafood wholesale market in Wuhan. An exception being one woman who had not been to the market but has close contact with her husband, one of the patients who worked in the market before falling sick, suggesting the possibility that the agent can at least undergo a limited degree of human to human transmission. Apart from this couple, there is no strong evidence which suggests that the unknown agent is highly infectious as no health care personnel in the hospitals where the patients were admitted were infected. Three suspicious cases were reported outside China, two in Thailand and one in Japan. These three patients were known to originated from or have been to Wuhan.

On 6^th^ January 2020, the Chinese authority released the sequence (accession#: MN908947) of a novel coronavirus, designated as 2019-nCoV, which was isolated from one of the pneumonia patients and confirmed to be the causative agent for this outbreak. Coronaviruses are a large family of viruses, most of which cause mild infections such as the common cold, but some such as the SARS and MERS (Middle East Respiratory Syndrome) viruses cause severe and potential fatal respiratory tract infections. Some coronaviruses are known to be transmitted easily between humans, while others do not. Based on currently available information, the 2019-nCoV virus belongs to a category that can cause severe illness in some patients but does not transmit readily between people. It is necessary to investigate the genetic and functional data of this new virus and compare to other coronaviruses so as to guide future research and design of appropriate infection control policy to prevent widespread dissemination of another potentially deadly coronavirus since the emergence of the SARS and MERS viruses. In this study, we performed in-depth genetic analysis of 2019-nCoV and generated data which provide timely and valuable insight into the potential origin of this virus, its ability to cause human infection, and its genetic relatedness with SARS and MERS.

Phylogenetic analysis of genomic sequences of coronaviruses deposited in the GenBank revealed that 2019-nCoV belonged to betacoronavirus and exhibited the closest linkage with two SARS-like coronavirus from bat (bat-SL-CoVZX45 and bat-SL-CoVZX21) (**Fig 1a**). According to the phylogenetic tree, human SARS viruses were closest to bat SARS-like viruses but with a lesser degree to bat coronaviruses, and was least related to other coronaviruses. The 2019-nCoV stands in a position between bat SARS-like viruses and bat coronaviruses, suggesting it is less related to the human SARS virus than other bat SARS-like viruses, and is likely a new type of bat coronavirus. Nevertheless, all coronaviruses that exhibit close linkage with 2019-nCoV originated from bat, strongly suggesting that this new coronavirus originated from bat (**Fig 1a**). Coronaviruses of other species including the murine coronavirus are genetically distant from this new coronavirus, indicating that 2019-nCoV did not originate from other animal hosts. As bats are not sold in the Wuhan market, animals that serve as the transmission vehicle remains to be identified (*1, 2*).

**Figure 1.**
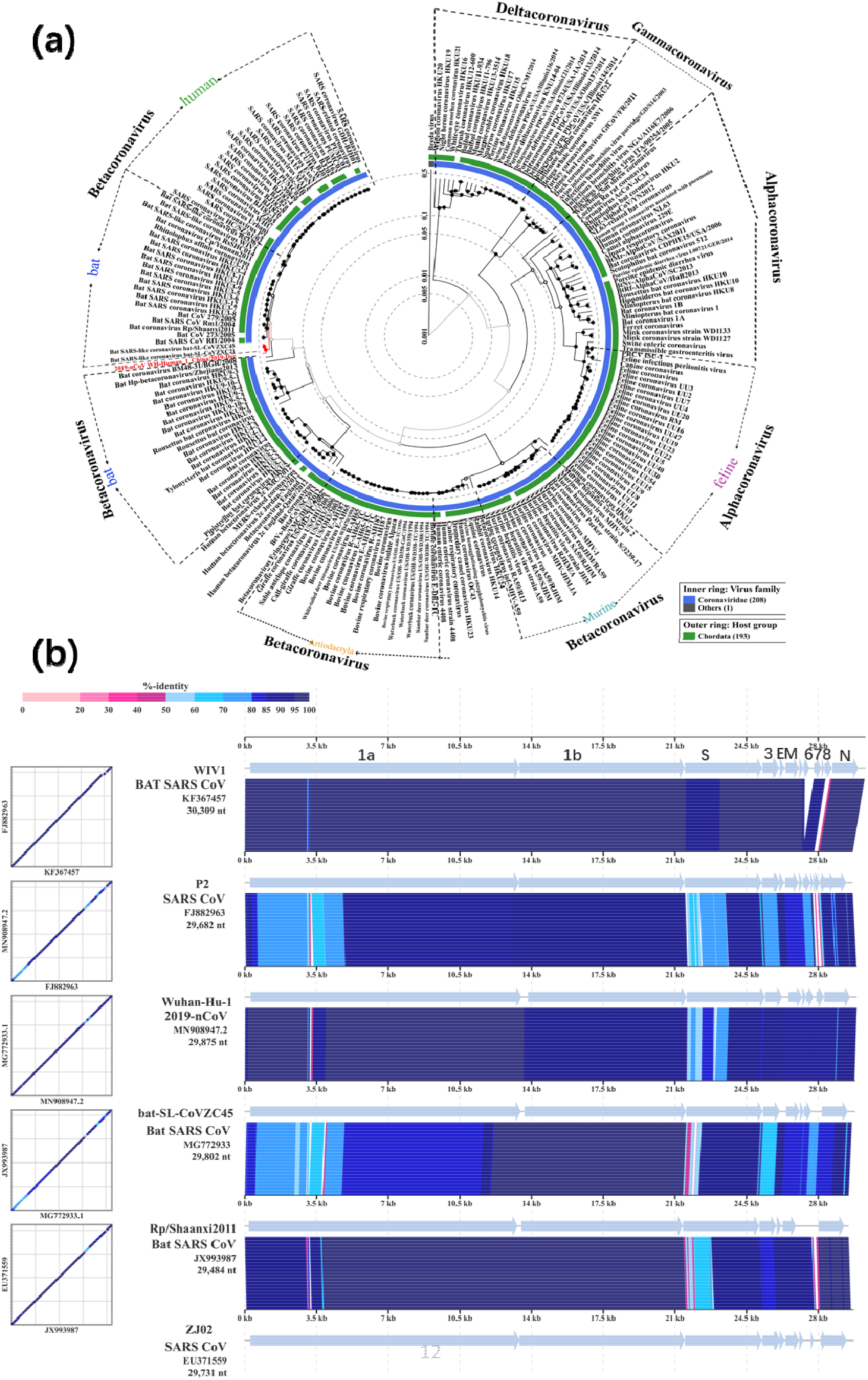
Phylogenetic analysis and sequence alignment of coronoviruses of different species. (a) Phylogenetic tree of coronaviruses from different species. The type of coronovirus and the host were labelled. Virus labeled with red is the newly discovered coronovirus 2019-nCoV. (b) Sequence alignment of representative the new 2019-nCoV, bat SARS like coronoviruses and bat coronoviruses. These included two highly homologous human SARS coronaviruses: SARS CoV P2 (FJ882963) and SARS CoV ZJ02 (EU371559), one bat SARS virus showing high homology with human SARS virus and similar potential to infect human as human SARS coronavirus: bat SARS CoV W1V1 (KF367457), two bat SARS-like viruses that are not able to infect human: (bat-SL-CoVZX45 and Rp Shaanxi2011) and the newly discovered 2019-nCoV from Wuhan.

The sequence of 2019-nCoV was annotated and aligned with several representative coronaviruses selected according to the degree of genetic relatedness depicted by the phylogenetic tree (**Fig 1b**). These included two highly homologous human SARS coronavirus: SARS CoV P2 (FJ882963) and SARS CoV ZJ02 (EU371559), one bat SARS virus that exhibits high homology with human SARS virus and similar potential to infect human as human SARS coronavirus: bat SARS CoV W1V1 (KF367457) (*3*), two bat SARS-like viruses that were not able to infect human: (bat-SL-CoVZX45 and Rp Shaanxi2011), and two un-related coronaviruses, the MERS virus MERS CoV (NC019843) and the Avian Infectious Bronchitis (IBV) virus IBV CoV (AY646283). The new 2019-nCoV was annotated slightly different from the human SARS virus and other coronaviruses, but the functionally important ORFs, ORF1a and ORF1b, and major structural proteins including the spike (S), membrane (M) and envelop (E) and nucleic capsid (N) proteins are well annotated (**Fig 1b**). Consistent with the phylogenetic tree data, 2019-nCoV did not align well with the MERS and IBV virus (**Fig S1**). Among the SARS viruses, bat coronaviruses and 2019-nCoV, the non-structural proteins generally aligned well but variations were observable in major structural proteins and some small ORFs (**Fig 1b**). Detailed sequence alignment showed that 2019-nCoV exhibited significant sequence variation at several regions with the human SARS coronavirus including the N-terminal region of ORF1a and S protein, ORF3, E, ORF6, 7 and 8, and the middle part of N protein. Bat SARS-like virus W1V1 aligned well with human SARS virus P2, with some variations at ORF8 and an insertion between ORF6 and 7. Coronavirus 2019-nCoV aligned best with the bat SARS-like virus bat-SL-CoVZX45, with the majority of genetic variations being seen at the N-terminal part of S protein. The bat-SL-CoVZX45 virus itself exhibited a high degree of variation with another bat SARS-like CoV Rp Shaanxi2011 at ORF1a, the N-terminus of S protein and other structural proteins. However, bat SARS-like CoV Rp Shaanxi2011 exhibited high homology with human SARS virus ZJ02, with variation being seen at the N-terminus of S protein and the middle part of N protein (**Fig 1b**). To check if 2019-nCoV is more close to bat SARS like virus or bat coronavirus, we selected two adjacent viruses from each group, bat SARSCoV Rf1/2004 and bat coronavirus BM48-31/BGR/2008, to perform alignment with the new virus with result showing that 2019-nCoV showed similar diversity from these two strains (**Fig S1b**). When 2019-nCoV was aligned with bat CoV HKU9-1, another bat coronavirus with further distance, it showed that 2019-nCoV was very different from this virus (**Fig S1c**). These sequence alignment data were consistent with results of phylogenetic tree analysis and indicated that 2019-nCoV exhibited stands in between bat SARS like viruses and bat coronaviruses, but is genetically distant from the human SARS virus. It should be considered a new type of bat coronavirus.

Since the S protein is the protein that exhibits the highest degree of genetic variations among different coronaviruses, we performed phylogenetic analysis of the S protein of different coronaviruses (**Fig S2**). Our data showed that the S protein of 2019-nCoV exhibited high homology with bat SARS-like coronaviruses such as bat-SL-CoVZX45 and bat-SL-CoVZX21, human SARS virus and bat coronaviruses. Homology of S protein of 2019-nCoV with representative human SARS virus, bat SARS like viruses and bat coronaviruses was determined as shown in Table S2. S protein of 2019-nCoV showed about 76% homology to human SARS virus P2 and high homology to bat SARS like viruses, while it showed 72% homology to closest bat coronavirus, bat coronavirus BM48-31, even lower homology with other bat coronaviruses. These data further suggested that 2019-nCoV is more likely a new type of bat coronavirus with only loose linkage with the SARS virus. Interestingly, different regions of the S proteins exhibited different levels of homology among the known coronaviruses (**Fig 1b**). Amino acid sequence alignment showed consistently that the N-terminal regions were far more diverse than the C-terminus, which seemed to be highly conservative (**Fig S3a**). Aligning these regions to the structure of S protein indicated that the structurally conserved C-terminal aligned well to the transmembrane domain which consists of a double helix, whereas the most variable region aligned to the N-terminal domain; on the other hand, the receptor binding domain exhibited intermediate level of sequence variation (**Fig S3b**).

Due to the high amino acid sequence homology of the S protein in 2019-nCoV and the SARS virus which can cause severe human infection, we analyzed the structural similarity of this protein in various viruses. Protein structure modeling was performed to obtain high quality structure of S proteins from different coronaviruses (**Table S1**). The high level similarity observable between the structures of S protein from different viruses implied that the S protein of 2019-nCoV and bat coronaviruses would most likely use the same human cell receptor as SARS virus. It was shown that angiotensin-converting enzyme 2 (ACE2) was the cellular receptor of the SARS virus S protein (*4*). Complex structure of ACE2 with the receptor binding domain (RBD) of S protein of SARS virus has been resolved and demonstrated tight interaction between these two proteins at the interaction interface (*5*). These data prompted us to determine the level of interaction between the S protein of 2019-nCoV with its potential cellular receptor ACE2. Using modeled S protein structure and by further performing structure-based alignment, we obtained the complex structure of RBD of the S protein of 2019-nCoV and several bat coronaviruses with human ACE2, using the complex structure of RBS of S protein from human SARS virus and ACE2 (2ajf) as reference (*5*). Structural analysis of potential interactions between RBD of S protein from human SARS virus and ACE2 protein depicted several interaction points including four hydrophobic interactions: ACE2(Y^41^)/ RBD(Y^484^), ACE2(L^45^)/ RBD(Y^484^), ACE2(L^79^, M^82^)/ RBD(L^472^), ACE2(Y^83^)/ RBD(Y^475^), one salt-bridge: ACE2(E^329^)/ RBD(R^426^) and one cation-π interaction: ACE2(K^353^)/ RBD(Y^491^) (**Fig 2a, 2e**). However, examination of interaction between RBD of 2019-nCoV and human ACE2 depicted only one potential hydrophobic interaction between ACE2(L^79^, M^82^) and RBD(F^486^) and one cation-π interaction interaction ACE2(K^353^)/ RBD(Y^492^). Further examination of interactions between RBD from bat SARS like coronaviruses, bat-SL-CoVZX45 and Rp Shaanxi2011 that do not infect human, showed only one cation-π interaction interaction, ACE2(K^353^)/ RBD(Y^481^). Another bat-originated coronavirus, bat SARS CoV W1V1 that displayed strong binding to ACE2 and exhibited potential to cause human infection was also included for analysis. Binding affinity of RBD from this virus to ACE2 was as tight as that of the human SARS virus, involving four hydrophobic interactions: ACE2(Y^41^)/ RBD(Y^485^), ACE2(L^45^)/ RBD(Y^485^), ACE2(L^79^, M^82^)/ RBD(F^473^), ACE2(Y^83^)/ RBD(Y^476^), one salt-bridge: ACE2(E^329^)/ RBD(R^427^) and one cation-π interaction site ACE2(K^353^)/ RBD(Y^492^). These data suggested that the higher binding affinity of RBD of coronavirus to ACE2 will confer the virus higher infectivity and pathogenicity. The fact that the RBD of 2019-nCoV exhibited much lower affinity to ACE2 implies that the virulence potential of 2019-nCoV should be much lower than that of human SARS virus, but is nevertheless stronger than viruses that do not cause human infection; such finding is also consistent with the current epidemiological data in that 2019-nCoV only caused severe pneumonia in patients with weaker immune system such as the elderlies and people with underlying diseases. The weaker binding affinity of 2019-nCoV to human cell might also explain the limited human to human transmission potential of this virus observed to date.

**Figure 2.**
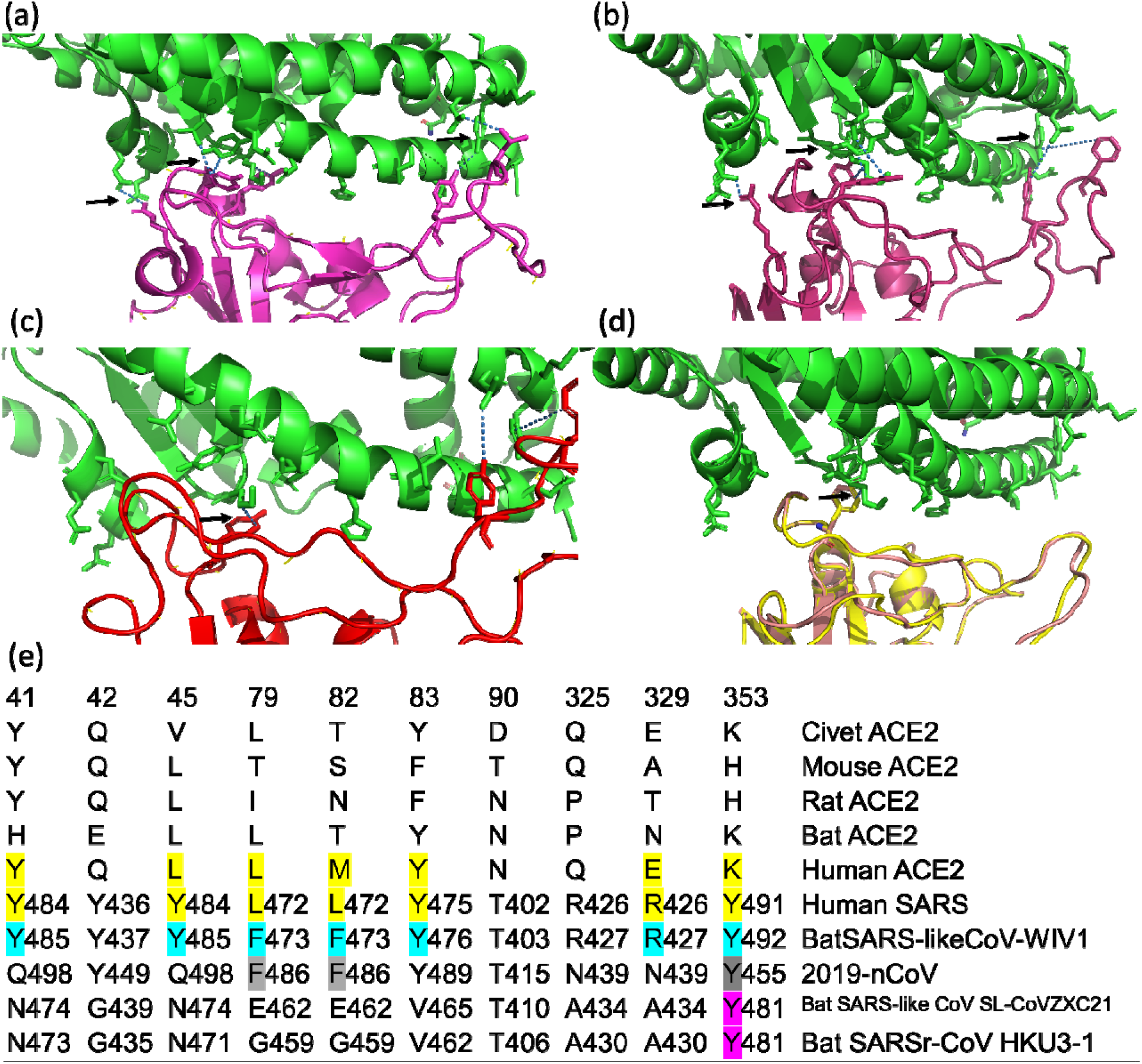
Potential interactions between receptor binding domain (RBD) of S proteins from different coronaviruses and the human cell receptor ACE2. Interactions between ACE2 with RBD of human SARS virus (a), highly similar bat SARS like coronavirus, CoV-W1V1 that can infect human (b), new type of coronavirus, 2019-nCoV (c), bat SARS like coronavirus CoV SL-CoVZXC21 and bat coronavirus HKU3-1 (d) are shown. The detailed amino acid interaction sites between these two proteins are shown in (e). Arrows showed the areas with interacted residues from both proteins. Amino acid highlighted with different colors indicated the potential interaction residues between different proteins, which was highlighted with different colors.

In this study, we utilized the whole genome sequence of the newly discovered coronavirus, 2019-nCoV, that caused an outbreak of pneumonia in Wuhan, China to perform comparative genetic and functional analysis with the human SARS virus and coronaviruses recovered from different animals. Phylogenetic analysis of coronavirus of different species indicated that 2019-nCoV might have originated from bat, but the intermediate transmission vehicle is not known at this stage. Genetic linkage analysis showed that 2019-nCoV lied at the interface between bat SARS like coronavirus and bat coronavirus and should belong to a novel type of bat coronavirus owing to high degree of variation from the human SARS virus. Analysis of the potential interaction of RBD of 2019-nCoV with human ACE2 receptor protein indicated that its affinity to human cell is much lower than that of human SARS virus due to the loss of several important interaction sites, implying that the infectivity and pathogenicity of this new virus should be much lower than the human SARS virus. These data facilitate design of appropriate policies to control further dissemination of this new coronavirus.

As of Jan 22, 2020, the infection cases were sharply increased in the past few days reaching a total of 440 cases with 9 confirmed death. Sporadic new cases were reported in more and more provinces in China. Human to human transmission has been confirmed as over 15 healthcare personnel have been confirmed to be infected. These epidemiological data were quite different from the data reported in the beginning and may suggest that the new virus could undergo human host adaption / evolution and become more adaptive to human host leading to more efficient human to human transmission. It is urgent to isolate and sequence the virus from the most recent cases to trace the evolutional mutations of the new virus. Using the analysis platform that we have developed above, we should be able to predict whether the new mutations could lead to the increase of infectivity of the mutated virus in a very short time.

## Acknowledgments

We acknowledge generation and provision of the genome sequence of 2019-nCoV (deposited in GenBank with the accession number MN908947) by Professor Yong-Zhen Zhang from The Shanghai Public Health Clinical Center & School of Public Health, and scientists from the partner institutions including the Central Hospital of Wuhan, Huazhong University of Science and Technology, the Wuhan Center for Disease Control and Prevention, the National Institute for Communicable Disease Control and Prevention, Chinese Center for Disease Control, and the University of Sydney, Sydney, Australia. We also appreciate contribution by authors who deposited the coronavirus sequences that we have used in this study in GenBank. This study has no intention to claim any credit of the whole genome sequence of 2019-nCoV and was designed with the only purpose of providing insight into the genetic and functional characteristics of this newly identified coronavirus through the utilization and analysis of sequences generated by the above acknowledged person and institutions.

## Conflicts of interest

We declare that we have no conflicts of interest.

## Reference

1. S. K. Lau et al., Severe acute respiratory syndrome coronavirus-like virus in Chinese horseshoe bats. Proc Natl Acad Sci U S A 102, 14040–14045 (2005).

2. K. K. To, I. F. Hung, J. F. Chan, K. Y. Yuen, From SARS coronavirus to novel animal and human coronaviruses. J Thorac Dis 5 Suppl 2, S103–108 (2013).

3. X. Y. Ge et al., Isolation and characterization of a bat SARS-like coronavirus that uses the ACE2 receptor. Nature 503, 535–538 (2013).

4. W. Li et al., Angiotensin-converting enzyme 2 is a functional receptor for the SARS coronavirus. Nature 426, 450–454 (2003).

5. F. Li, W. Li, M. Farzan, S. C. Harrison, Structure of SARS coronavirus spike receptor-binding domain complexed with receptor. Science 309, 1864–1868 (2005).

