## Supplementary method, tables and figures for "Genomic and protein structure modelling analysis depicts the origin and infectivity of 2019-nCoV, a new coronavirus which caused a pneumonia outbreak in Wuhan, China"

#contribute equally to the work.

*Corresponding author: Sheng Chen, City University of Hong Kong, Kowloon, Hong Kong;

**Keywords:** 2019-nCoV, Genomics, Protein modelling, Origin, Infectivity, Wuhan

**Supplementary methods**

**Phylogenetic analysis and sequence alignment**

A total of 211 genome sequences of viruses in the Coronavirinae family including the 2019-nCoV Wuhan virus were downloaded from GenBank (last accessed on 11 January 2020). Circular proteomic trees were computed using ViPTreeGen v1.1.2 (*1*). The sequence of a Breda virus (accession: NC_007447) was used as an outgroup. Alignment of sequences in different viral genomes was conducted using the alignment function of ViPTree (*1*). The phylogenetic tree and sequence alignment products were manually edited using Inkscape v0.91 (*2*). Ten spike protein sequences which were similar to that of 2019nCoV were downloaded from NCBI. SNP analysis was performed using MegaX(*3*), and the alignment was carried out by using ClustalW(*4*). The aligned sequences were edited and viewed in CLC Genomics Workbench 20.

**Protein structure prediction and contacts between the human ACE2 and spike receptor-binding domains**

Spike receptor-binding domain (RBD) of coronavirus 2019-nCoV and four bat-originated coronavirus were predicted by aligning their spike protein sequences to spike RBD of SARS coronavirus (*5*) using Clustal Omega (*6*). Homology modeling of spike proteins and the related RBDs of 2019-nCoV and four bat-originated coronavirus was performed using SARS-CoV spike glycoprotein as template (PDB ID: 5X58) (*7*) on the Swiss-Model workspace (*8*). The structure assessment results are presented below. The models were visualized by PyMol. The contacts between human ACE2 and spike RBD were predicted by aligning to structure of the RBD in complex with the human receptor ACE2 (PDB ID: 2AJF) (*5*).

**Supplementary Table S1. Assessment of the quality of modeled structures of spike protein and its receptor binding domain (RBD) of different coronavirus using protein structures of the human SARS virus as templates.**

| **S protein** | **GenBank** | **residues** | **MolProbity Score^1^** | **Ramachandran Favoured^1^** | **GMQE^2^** | **QMEAN^2^** |
| --- | --- | --- | --- | --- | --- | --- |
| 2019-nCoV | MN908947 | 1273 | 1.64 | 89.4% | 0.73 | -4.46 |
| BatSARS-likeCoV-WIV1 | KC881007 | 1256 | 1.36 | 90.02% | 0.74 | -3.45 |
| BatSARS-likeCoV | MG772934 | 1245 | 1.64 | 88.96% | 0.73 | -3.91 |
| BatSARSr-CoV | DQ022305 | 1242 | 1.34 | 89.49% | 0.73 | -3.96 |
| BatCoV | JX993987 | 1240 | 1.4 | 90.02% | 0.73 | -3.90 |

1, Structure assessment. MolProbity is all-atom contact analysis based only on properties of the predicted model. Lower numbers indicate better models. Ramachandran favoured indicates energetically favoured regions for backbone dihedral angles against amino acid residues in protein structure. Larger numbers indicate better models.

2, Model evaluation. GMQE (Global Model Quality Estimation) is a quality estimation which combines properties of the target–template alignment and the template search method. The resulting GMQE score is expressed as a number between 0 and 1, larger numbers indicate higher reliability. The QMEAN Z-score provides an estimate of the "degree of nativeness" of the structural features observed in the model on a global scale. QMEAN Z-scores around zero indicate good agreement between the model structure and experimental structures of similar size.

**Supplementary Table S2. Homology of S protein between different coronaviruses.**

|  | **2019-nCoV** | **SL-CoVZXC21** | **CoV-HKU3-1** | **CoV-WIV1** | **SARS_P2** | **Shaanxi-2011** | **Bat-BM48-31** | **Bat-zj2013** | **CoV-HKU5-1** | **CoV-HKU4-1** |
| --- | --- | --- | --- | --- | --- | --- | --- | --- | --- | --- |
| **2019-nCoV** | 100% | 81.35% | 77.93% | 77.7% | 76.58% | 76.56% | 72.5% | 41.06% | 31.21% | 30.58% |
| **SLCoVZXC21** | 81.35% | 100% | 83.09% | 78.15% | 77.5% | 82.05% | 71.8% | 40.59% | 31.62% | 31.46% |
| **CoV-HKU3-1** | 77.93% | 83.09% | 100% | 80.11% | 79.55% | 88.38% | 75.1% | 40.79% | 32.27% | 31.87% |
| **CoV-WIV1** | 77.7% | 78.15% | 80.11% | 100% | 92.11% | 81.06% | 75.66% | 41.35% | 31.31% | 30.8% |
| **SARS_P2** | 76.58% | 77.5% | 79.55% | 92.11% | 100% | 81.14% | 75.64% | 41.14% | 31.42% | 30.99% |
| **Shaanxi-2011** | 76.56% | 82.05% | 88.38% | 81.06% | 81.14% | 100% | 75.06% | 41.6% | 31.91% | 31.67% |
| **Bat-BM48-31** | 72.5% | 71.8% | 75.1% | 75.66% | 75.64% | 75.06% | 100% | 40.21% | 32.12% | 32.05% |
| **Bat-zj2013** | 41.06% | 40.59% | 40.79% | 41.35% | 41.14% | 41.6% | 40.21% | 100% | 28.77% | 28.9% |
| **CoV-HKU5-1** | 31.21% | 31.62% | 32.27% | 31.31% | 31.42% | 31.91% | 32.12% | 28.77% | 100% | 69.23% |
| **CoV-HKU4-1** | 30.58% | 31.46% | 31.87% | 30.8% | 30.99% | 31.67% | 32.05% | 28.9% | 69.23% | 100% |

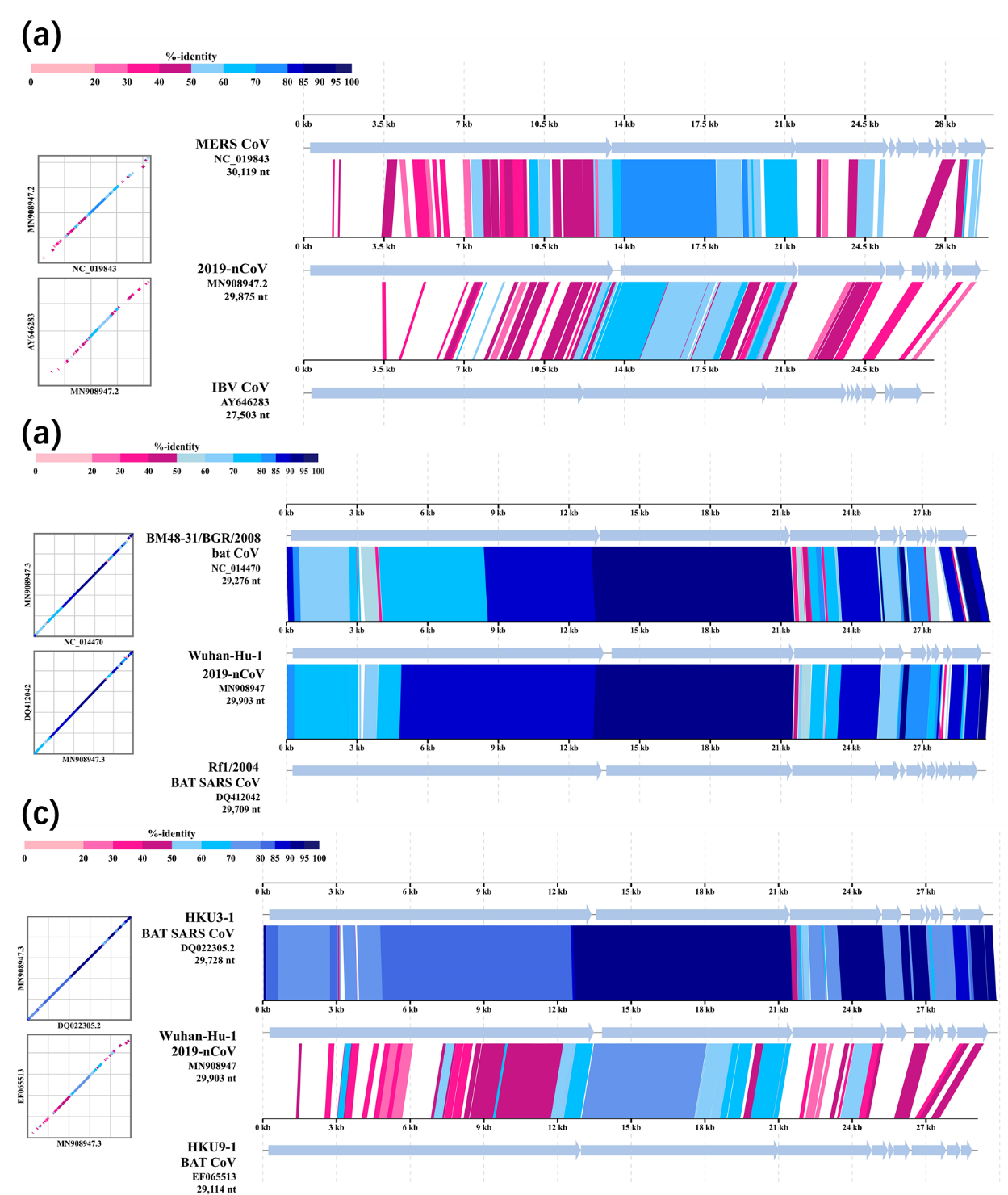

**Supplementary Figure S1. Sequence alignment of 2019-nCoV with Middle East Respiratory Syndrome (MERS) virus: MERS CoV (NC019843) and Avian Infectious Bronchitis (IBV) virus: IBV CoV (AY646283).** The new coronavirus showed very low homology with these two types of coronaviruses.

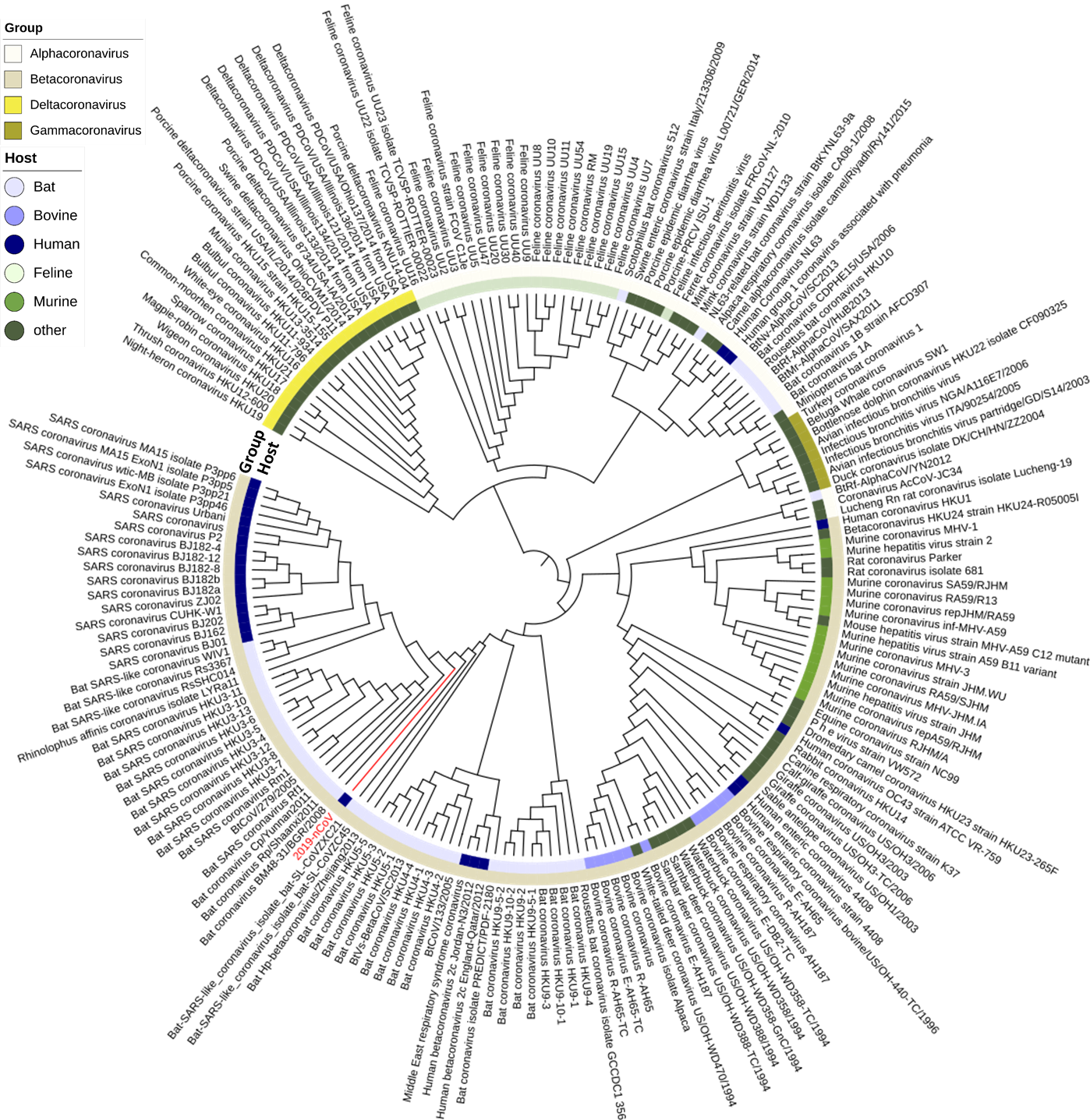

**Supplementary Figure S2. Phylogenetic analysis of the spike (S) protein of different coronaviruses.** The coronaviruses selected for this analysis are the same as in Figure 1a. The pattern of genetic linkage among sequences of the S protein of various viruses is slightly different from that of the whole genome sequences. In particular, the S protein of 2019-nCoV is at a position further away from the human SARS coronavirus, yet the protein is closer to bat SARS-like coronaviruses than other bat coronaviruses. Different coronavirus groups and the respective hosts are labelled.

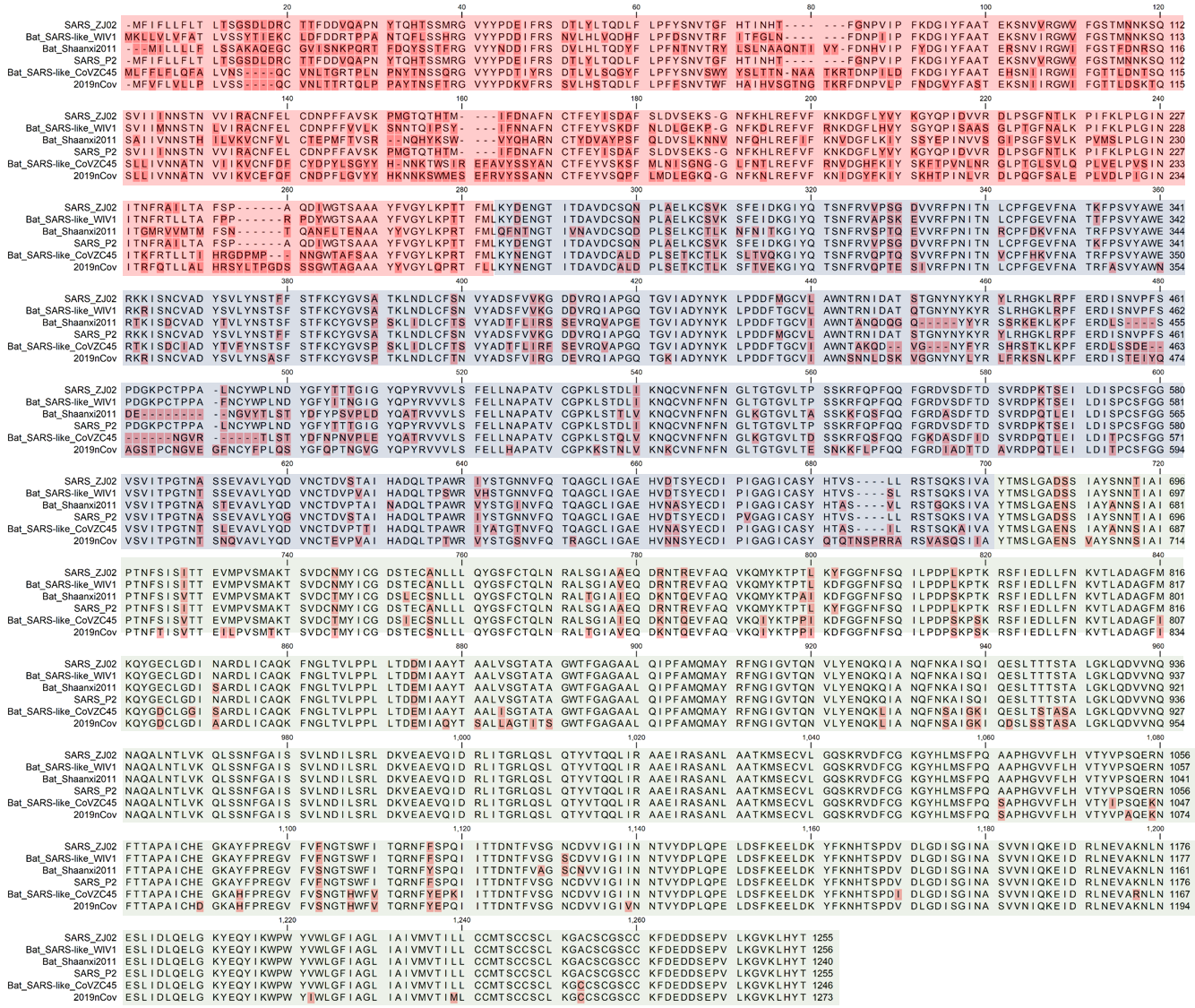

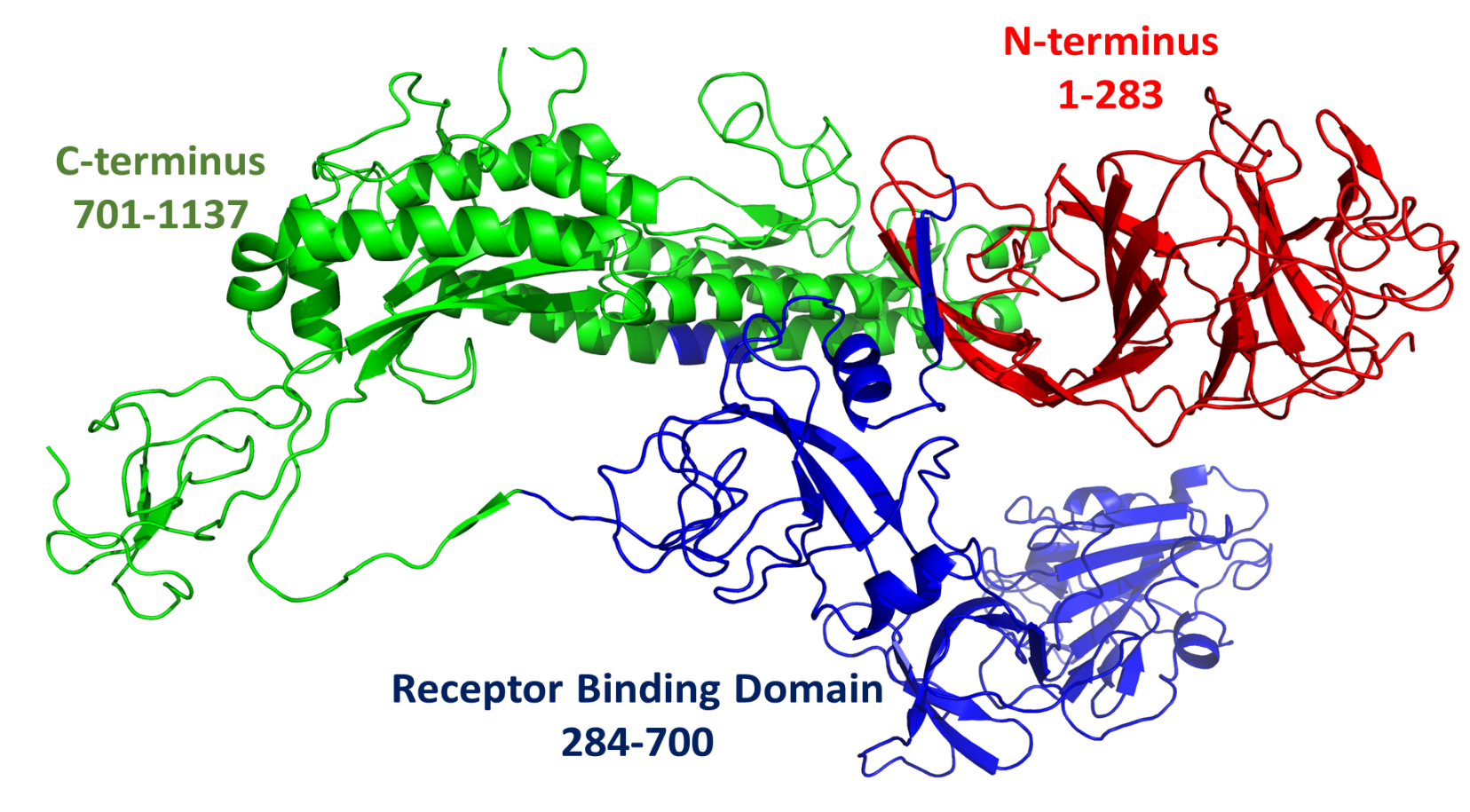

**Supplementary Figure S3. Amino acid sequence alignment of coronaviruses from bat and human.** Upper panel is the amino acid sequence alignment and the lower panel is the structure of the S protein. Three domains of S protein were labelled as red, blue and green, which aligned with the different colors for the aligned sequence. Sequences that exhibits the highest degree of variation are in the N-terminal domain, followed by those in the RBD. The C-terminal green domain is the most conservative among the test viruses.

**References:**

1.	Y. Nishimura *et al.*, ViPTree: the viral proteomic tree server. *Bioinformatics* **33**, 2379-2380 (2017).

2.	B. T., Inkscape: guide to a vector drawing program[M]. *Upper Saddle River, NJ, USA: Prentice Hall,* , (2011).

3.	S. Kumar, G. Stecher, M. Li, C. Knyaz, K. Tamura, MEGA X: molecular evolutionary genetics analysis across computing platforms. *Molecular biology and evolution* **35**, 1547-1549 (2018).

4.	M. A. Larkin *et al.*, Clustal W and Clustal X version 2.0. *Bioinformatics* **23**, 2947-2948 (2007).

5.	F. Li, W. Li, M. Farzan, S. C. Harrison, Structure of SARS coronavirus spike receptor-binding domain complexed with receptor. *Science* **309**, 1864-1868 (2005).

6.	F. Madeira *et al.*, The EMBL-EBI search and sequence analysis tools APIs in 2019. *Nucleic acids research* **47**, W636-W641 (2019).

7.	Y. Yuan *et al.*, Cryo-EM structures of MERS-CoV and SARS-CoV spike glycoproteins reveal the dynamic receptor binding domains. *Nature Communications* **8**, 15092 (2017).

8.	A. Waterhouse *et al.*, SWISS-MODEL: homology modelling of protein structures and complexes. *Nucleic Acids Research* **46**, W296-W303 (2018).
